# Cross-Species Protection to Innate Immunity Mediated by A Bacterial Pigment

**DOI:** 10.1101/2023.01.15.524085

**Authors:** Yiwei Liu, Eleanor A. McQuillen, Pranav S. J. B. Rana, Erin. S. Gloag, Daniel J. Wozniak

## Abstract

Bacterial infections are often polymicrobial. *Pseudomonas aeruginosa* and *Staphylococcus aureus* cause chronic co-infections, which are more problematic than mono-species infections. We found that the production of *S. aureus* membrane-bound pigment staphyloxanthin (STX), was induced by the *P. aeruginosa* exoproduct, 2-heptyl-4-hydroxyquinoline N-oxide (HQNO). The induction phenotype was conserved in *P. aeruginosa* and *S. aureus* clinical isolates examined. When subjected to hydrogen peroxide or human neutrophils, *P. aeruginosa* survival was significantly higher when mixed with wild-type (WT) *S. aureus*, compared to a mutant deficient in STX production or *P. aeruginosa* alone. In a murine wound model, co-infection with WT *S. aureus*, but not the STX-deficient mutant, enhanced *P. aeruginosa* burden and disease compared to mono-infection. In conclusion, we discovered a novel role for *P. aeruginosa* HQNO mediating polymicrobial interactions with *S. aureus* by inducing STX production, which consequently promotes resistance of both pathogens to innate immune effectors. These results further our understanding of how different bacterial species cooperatively cause co-infections.

## INTRODUCTION

*Pseudomonas aeruginosa* and *Staphylococcus aureus* are two common microorganisms colonizing cystic fibrosis (CF) airways and chronic wounds^1-4^. Co-infection correlates with increased disease severity, compared to mono-infections caused by either species^4-6^.

*P. aeruginosa* and *S. aureus* have an intriguingly complicated relationship and have been used as model organisms to investigate polymicrobial interactions. *In vitro, P. aeruginosa* outcompetes *S. aureus*. Many antagonistic mechanisms have been determined in *P. aeruginosa*^7-10^. However, of relevance to this study is the *P. aeruginosa* quorum-sensing (QS) system PQS (*Pseudomonas* quinolone signal) which is crucial for antagonizing *S. aureus*^11^. The PQS system produces 2-heptyl-4-hydroxyquinoline n-oxide (HQNO) which inhibits respiration and promotes small colony variants formation in *S. aureus*^12^. In contrast to this antagonism *in vitro*, both pathogens can co-exist *in vivo*. In fact, interaction with *S. aureus* can benefit *P. aeruginosa* by increased biofilm formation^13^, host immunity evasion^14^, and antibiotic resistance^15,16^. One of the keys to understanding this polymicrobial relationship is the subtle balance between the competitive and cooperative behaviors of these two organisms.

Staphyloxanthin (STX) is a membrane-bound yellow pigment, synthesized by the *crt* operon in *S. aureus*^17,18^ and widely produced among clinical and environmental isolates^19^. Strains deficient in STX production appear as white colonies on solid media. By functioning as an antioxidant to resist oxidative stress and altering membrane fluidity to combat antimicrobial peptides (APs), STX mediates *S. aureus* resistance to host defense mechanisms ^20,21^. Interestingly, a *P. aeruginosa* wound isolate was observed to induce STX production in a co-isolated white variant of *S. aureus*^22^. This implies that STX mediates interactions between *P. aeruginosa* and *S. aureus in vivo*.

Given the importance of *P. aeruginosa* and *S. aureus* co-infections and the implication of STX in polymicrobial interactions, here we investigated the role of STX during *P. aeruginosa* and *S. aureus* co-infections. We found that STX production is induced by *P. aeruginosa* HQNO and affords cross-species protection against host innate immunity.

## RESULTS

### Both surface- and planktonic-grown *P. aeruginosa* can induce *S. aureus* STX production

In our previous study, we examined the contribution of *P. aeruginosa* factors, especially released exopolysaccharide Psl^10^, in antagonizing the growth of *S. aureus* in a macrocolony proximity assay. *S. aureus* USA300 was grown at increasing distances from *P. aeruginosa* PAO1 on solidified media (Figure 1A). As expected, the growth of USA300 adjacent to PAO1 was inhibited (Figure 1A, black arrow). Interestingly, we also observed a difference in USA300 pigmentation. USA300 macrocolonies furthest from PAO1 were light yellow (Figure 1A, yellow arrows). However, the macrocolonies closer to PAO1 were more pigmented (Figure 1A, orange arrows), suggestive of elevated STX production. In contrast, macrocolonies of an *S. aureus crtM* transposon mutant (*crtM*::Tn, SAUSA300_2499) deficient in STX production, remained non-pigmented regardless of the distance to PAO1 macrocolonies (Figure 1B, white arrow). This suggests that an exoproduct secreted by PAO1 can diffuse through the solidified media to induce STX production in the nearby USA300.

**Figure 1.**
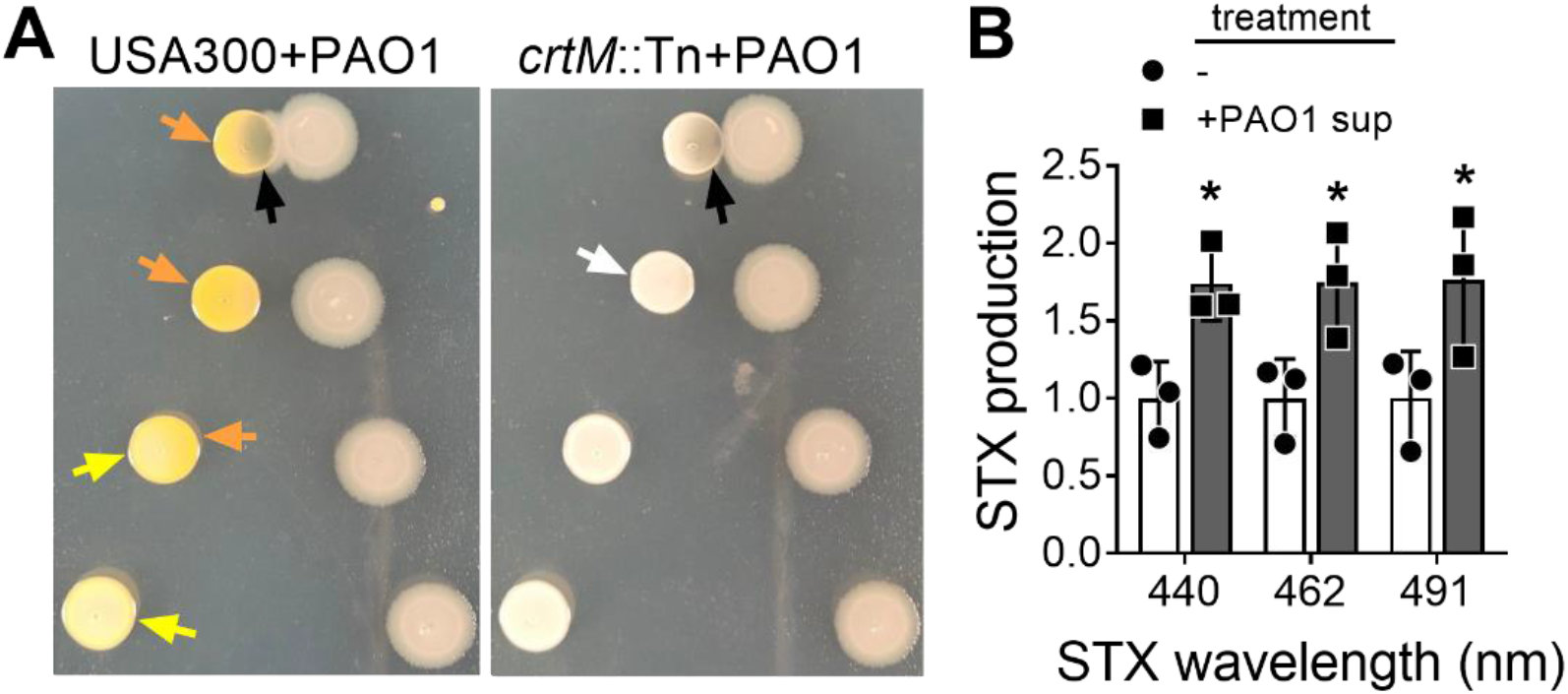
*S. aureus* STX production is induced by *P. aeruginosa* exoproduct(s). **(A)** USA300 (left) or *crtM*::Tn (right) was grown at increasing distances from PAO1 on solidified media in a macrocolony proximity assay. Yellow arrows point to USA300 with no pigment change, orange arrows point to USA300 with increased yellow pigmentation, and the white arrow indicates white *crtM*::Tn colonies. The black arrows point to *S. aureus* growth inhibition by PAO1. **(B)** STX production in USA300 treated with (+) or without (-) 5% filter-sterilized PAO1 spent media, was measured after methanol extraction at the wavelengths 440nm, 462nm, and 491nm. The results were normalized to the untreated group. Data are presented as mean ± SD from the results of 3 biological replicates, each with 2 technical replicates. *, *P*<0.05, compared to the untreated group.

STX production is a stress response of *S. aureus*^23^. Since there was increased STX production along with growth inhibition of USA300 when adjacent to PAO1 (Figure 1A), we investigated if STX induction by *P. aeruginosa* was a response of *S. aureus* to general growth inhibition. *S. aureus* macrocolonies were grown at increasing distances from either filter disks soaked in ciprofloxacin or daptomycin antibiotics or a PAO1 macrocolony (Figure S1). USA300 showed similar levels of growth inhibition by either antibiotics or PAO1 (black arrows). However, STX induction was only observed for USA300 macrocolonies grown in proximity to PAO1 (Figure S1C, orange arrow). Neither antibiotic induced STX production in USA300 macrocolonies (Figure S1AB). This implies that STX is induced in *S. aureus* in response to secreted *P. aeruginosa* factors and not a general response to growth inhibition.

Since exoproducts are secreted into the spent media during bacterial planktonic growth, we also examined if *S. aureus* STX could be induced by *P. aeruginosa* spent media when grown in planktonic culture. 50% PAO1 spent media demonstrated a strong bactericidal effect towards *S. aureus* within 4h of treatment^10^. To eliminate complications due to potential growth inhibition, we grew USA300 planktonic culture in 5% PAO1 spent media for 16h. STX was then extracted and quantified as described^17,18^. Indeed, PAO1 spent media induced *S. aureus* STX production (Figure 1C). The above data suggest that *P. aeruginosa* exoproduct(s) can induce STX production in WT *S. aureus*, regardless of the mode of growth.

### *P. aeruginosa* exoproduct HQNO induces *S. aureus* STX production

*P. aeruginosa* can secret many antagonistic factors, including Psl^10^, rhamnolipid^9^, HQNO^12^, LasA^7^, and pyoverdine^8^, to inhibit the growth of *S. aureus*. We used the corresponding mutants in PAO1 to determine if any of these mechanisms were responsible for inducing *S. aureus* STX production. When USA300 and these PAO1 mutants were grown together in the macrocolony proximity assay, Δ*pqsA* was the only mutant that neither antagonized nor induced STX production (Figure 2A), while others still antagonized the growth of USA300 and induced STX production to varying degrees (Figure S2). *pqsA* is the first gene in the *P. aeruginosa pqs* operon responsible for synthesizing two main end products, HQNO and PQS (3,4-dihydroxy-2-heptylquinoline)^24-26^. To investigate if either or both products were sufficient for STX induction, USA300 macrocolonies were grown at increasing distances from commercially acquired HQNO and PQS molecules, the solvent used to dissolve the chemicals, or PAO1 (Figure 2B). Increased STX production of USA300 macrocolonies was only observed when grown adjacent to HQNO and PAO1. Neither PQS nor the solvent induced STX production.

**Figure 2.**
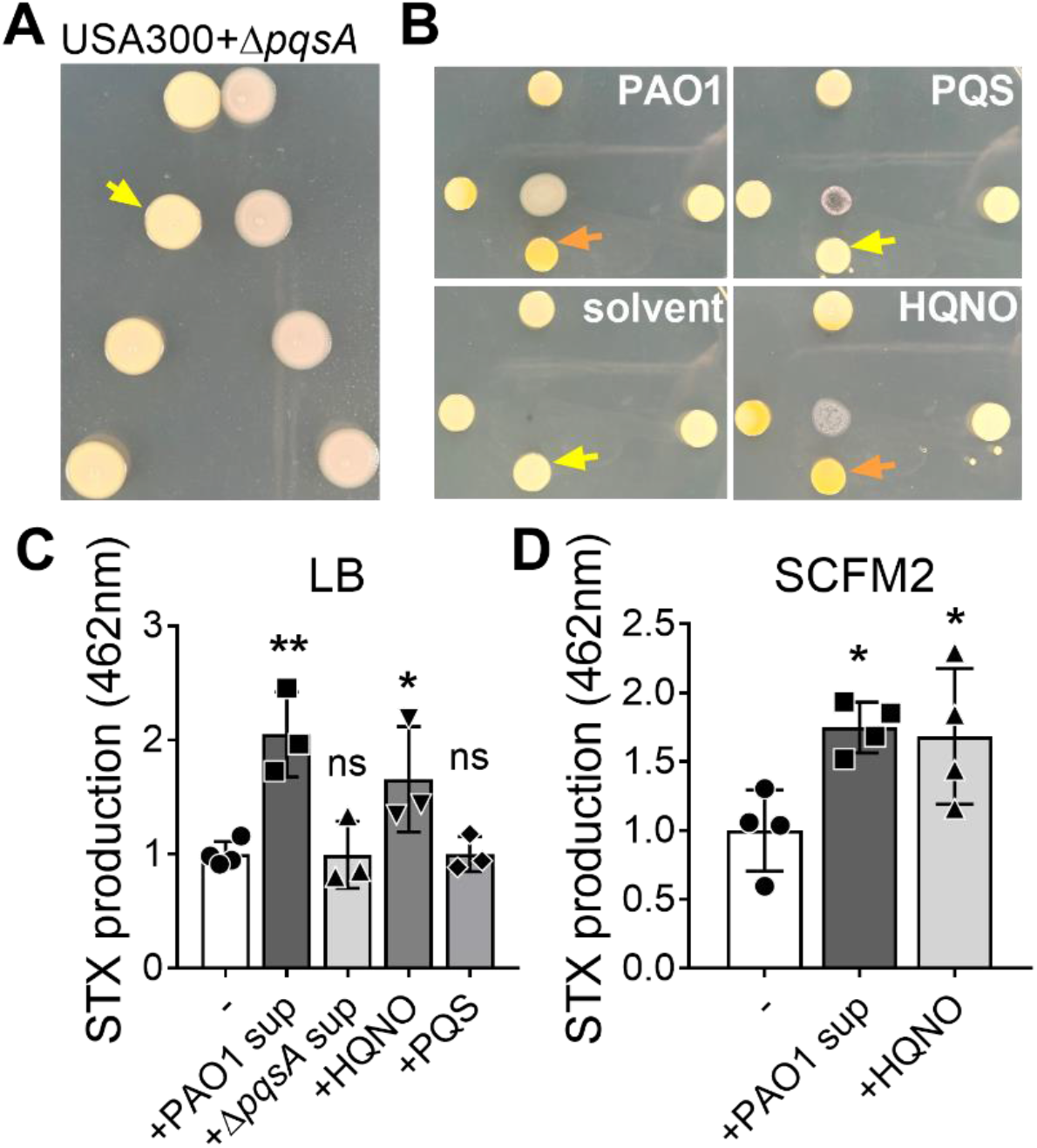
*P. aeruginosa* HQNO induces *S. aureus* STX production. USA300 was grown at increasing distances from **(A)** Δ*pqsA* and **(B)** PAO1, PQS, HQNO, or the solvent in which the molecules were dissolved, on solidified media in a macrocolony proximity assay. Yellow arrows point to USA300 colonies with no pigment change and orange arrows depict USA300 colonies with increased yellow pigmentation. **(C)** STX production in LB-grown USA300 treated with (+) or without (-) 5% filter-sterilized PAO1 or Δ*pqsA* spent media (sup), or 5μM of HQNO or PQS. STX production was quantified by absorbance at OD 462nm. **(D)** STX production in SCFM2-grown USA300 treated with or without 5% filter-sterilized PAO1 spent media (sup) or 5μM of HQNO was measured. For C and D, the results were normalized to the untreated group. Data are presented as mean ± SD from the results of at least 3 biological replicates, each with 2 technical replicates. *, *P*<0.05; **, *P*<0.01; ns, not significant, compared to the untreated group.

In addition to surface-grown *S. aureus*, we also examined if HQNO could induce STX production in planktonic *S. aureus* cultures. Since high concentrations of HQNO (e.g., 400μM) inhibit the growth of *S. aureus*^*27*^, we used a low concentration of 5μM. When grown in LB supplemented with either 5% PAO1 spent media or 5μM HQNO, USA300 produced significantly more STX compared to when grown in LB alone (Figure 2C). There was no significant difference in STX production between *S. aureus* grown in LB alone, or in media supplemented with PAO1Δ*pqsA* spent media or PQS (Figure 2C), indicating that neither was able to induce *S. aureus* STX production during planktonic growth.

Synthetic CF sputum media (SCFM2) mimics the CF sputum composition and has been used to culture both *P. aeruginosa* and *S. aureus*^28,29^. To determine if *P. aeruginosa* can induce STX production under *in vivo-*like conditions, USA300 was grown planktonically in SCFM2, supplemented with either PAO1 spent media or HQNO. In both conditions, STX production was significantly higher, compared to USA300 grown in SCFM2 alone (Figure 2D). Overall, the above data indicate that *P. aeruginosa* HQNO is sufficient to induce *S. aureus* STX production.

### *P. aeruginosa* induction of *S. aureus* STX production is prevalent among clinical isolates

Since STX production varies across *S. aureus* laboratory strains and clinical isolates^30,31^, we screened a collection of 61 *S. aureus* clinical isolates, from CF lung and bloodstream infections (Table S1) to examine their STX production in the presence of PAO1. Using the macrocolony proximity assay, *S. aureus* isolates were categorized into 3 classes based on colony pigmentation (Figure 3A, Table 1). 14.7% of *S. aureus* isolates (class I) remained white regardless of the distance to PAO1, suggestive of no STX production. 78.7% of isolates (class II) showed increased yellow pigmentation when grown in proximity to PAO1, suggesting STX induction. A small subset of these isolates appeared white when grown alone or distant from PAO1, similar to the phenotype reported by Antonic *et al*^22^. 6.6% of isolates (class III) had yellow macrocolonies, but the pigmentation did not change when grown adjacent to PAO1, suggesting that STX production was not further induced by *P. aeruginosa*. No significant difference was found between isolates derived from CF lungs or bloodstream infections. In addition to *S. aureus* clinical isolates, we also found that PAO1 could induce STX production in methicillin-sensitive *S. aureus* (MSSA) (Figure S3). Our screening indicates that STX induction by *P. aeruginosa* is prevalent among *S. aureus* clinical isolates.

**Table 1.** STX production and induction in *S. aureus* clinical isolates.

|  | Total #(% ) | I | II | III |
| --- | --- | --- | --- | --- |
| CF | 31 (100%) | 5 (16.1%) | 23 (74.2%) | 3 (9.7%) |
| Blood | 30 (100%) | 4 (13.3%) | 25 (83.3%) | 1 (3.3%) |
| Total | 61 (100%) | 9 (14.7%) | 48 (78.7%) | 4 (6.6%) |

I: no STX production
II: STX induced by PAO1
III: produces STX but no induction by PAO1
\* No significant difference was found when comparing the classifications of isolates derived from different sources.

**Figure 3.**
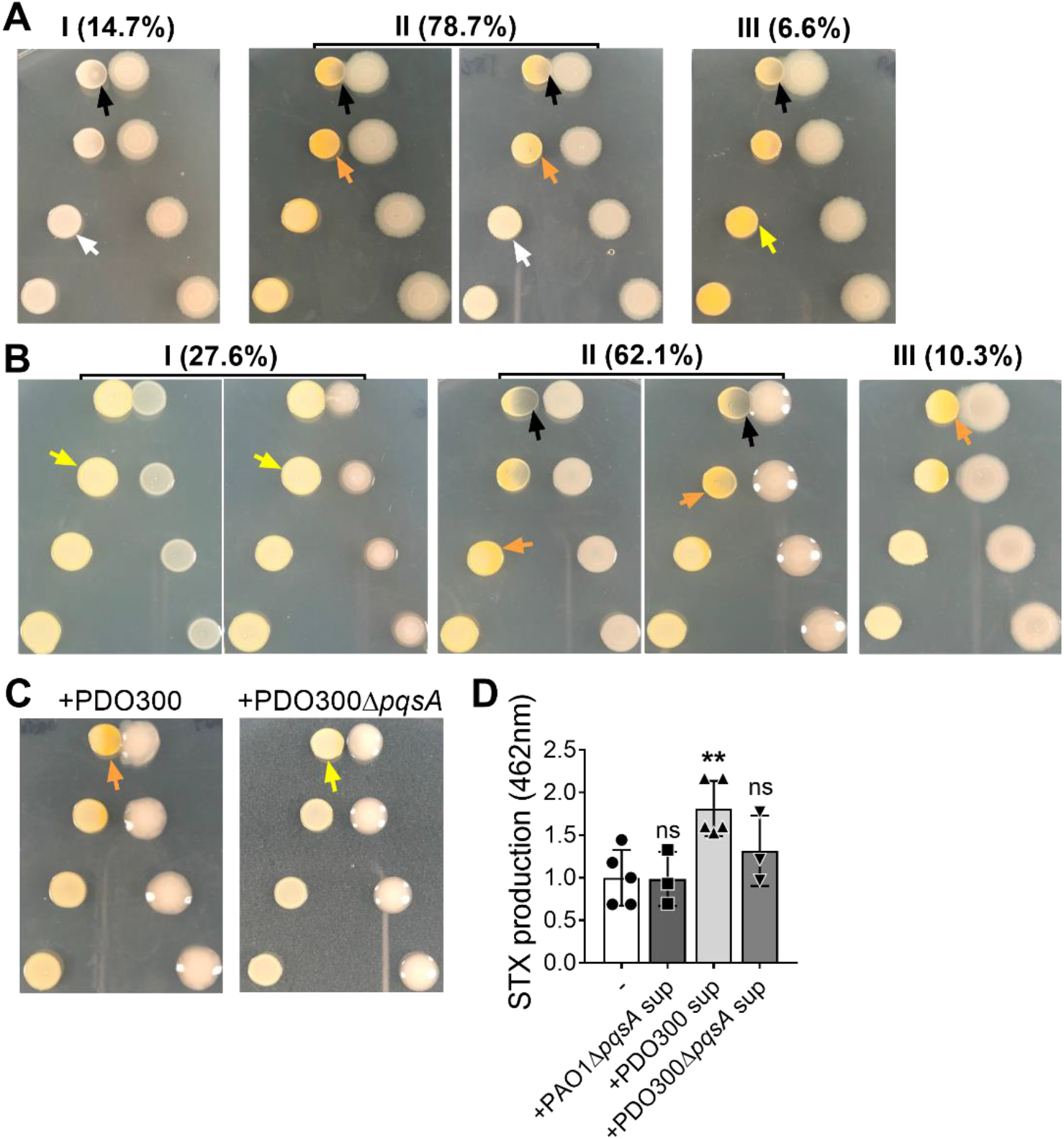
*P. aeruginosa* induction of *S. aureus* STX production is prevalent among clinical isolates. **(A)** Representative images and their respective proportions (%) of 3 classes of *S. aureus* clinical isolates when grown with *P. aeruginosa* PAO1 in a macrocolony proximity assay. Class I (14.7%) isolates were white colonies. STX production in Class II (78.7%), despite different intrinsic colors of the colonies (yellow: left; white: right), was induced by adjacent PAO1. Class III (6.6%) had yellow colonies, but no STX induction by PAO1. **(B)** Representative images and their respective proportions (%) of 3 classes of *P. aeruginosa* clinical isolates when grown with *S. aureus* USA300 in a macrocolony proximity assay. Class I (27.6%) did not induce STX production in the adjacent USA300 macrocolonies. Class II (62.1%) inhibited USA300 growth and induced STX production. Class III (10.3%) induced STX production in USA300 without growth inhibition. Both mucoid (right) and non-mucoid (left) strains were found in Class I and II. **(C)** USA300 was grown at different distances to mucoid PDO300 (left) or PDO300Δ*pqsA* (right) on solidified media in a macrocolony proximity assay. For A, B and C, the yellow arrows point to *S. aureus* colonies with no color change while the orange arrows depict *S. aureus* colonies with increased yellow pigmentation. The black arrows point to *S. aureus* growth inhibition by *P. aeruginosa*. **(D)** STX production in USA300, treated with (+) or without (-) 20% filter-sterilized PAO1Δ*pqsA*, PDO300 or PDO300Δ*pqsA* spent media (sup), was measured after methanol extraction at 462nm. The results were normalized to the untreated group. Data are presented as mean ± SD from the results of at least 3 biological replicates, each with 2 technical replicates. **, *P* < 0.01; ns, not significant, compared to the untreated group.

Additionally, *P. aeruginosa* clinical isolates synthesize varying levels of HQNO^32,33^. We screened 29 *P. aeruginosa* clinical isolates, derived from CF lung and wound infections (Table S1), to examine their ability to induce STX production in USA300. Using the macrocolony proximity assay, *P. aeruginosa* clinical isolates were also categorized into 3 classes (Figure 3B, Table 2). 72.4% of them were able to induce STX production. However, this group was further differentiated on the ability to inhibit the growth of USA300. Class II clinical isolates inhibited the growth of USA300 (62.1%), while class III did not (10.3%), suggesting that STX induction is independent of growth inhibition. The remaining 27.6% of *P. aeruginosa* clinical isolates did not induce STX production or inhibit the growth of USA300 (class I). No significant difference was found between isolates derived from CF lungs or wound infections. The above data suggest that the ability to induce *S. aureus* STX production is prevalent among *P. aeruginosa* clinical isolates.

**Table 2.**
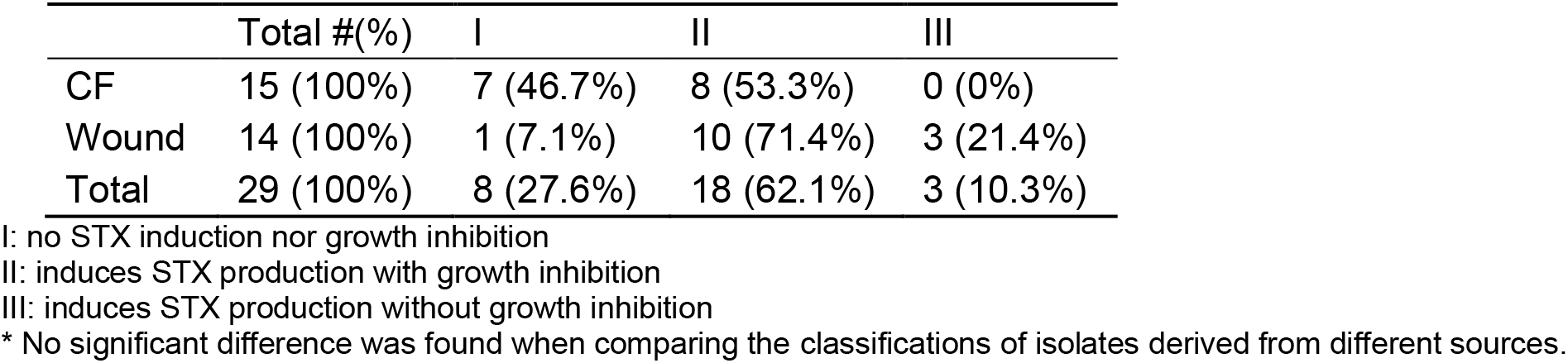
STX induction by *P. aeruginosa* clinical isolates.

|  | Total #(% ) | I | II | III |
| --- | --- | --- | --- | --- |
| CF | 15 (100%) | 7 (46.7%) | 8 (53.3%) | 0 (0%) |
| Wound | 14 (100%) | 1 (7.1%) | 10 (71.4%) | 3 (21.4%) |
| Total | 29 (100%) | 8 (27.6%) | 18 (62.1%) | 3 (10.3%) |

I: no STX induction nor growth inhibition
II: induces STX production with growth inhibition
III: induces STX production without growth inhibition
\* No significant difference was found when comparing the classifications of isolates derived from different sources.

Mucoid conversion, defined by overproduction of the exopolysaccharide alginate, occurs frequently in *P. aeruginosa* clinical strains^34^. Mucoid *P. aeruginosa* can co-exist with *S. aureus* better than non-mucoid counterparts, due to reduced production of antagonistic factors, including HQNO^35^. Since some of the *P. aeruginosa* clinical isolates tested above were mucoid but induced USA300 STX production (Figure 3B), we examined if mucoid *P. aeruginosa* could induce STX production in an HQNO-dependent manner. To test this, we used a laboratory *P. aeruginosa* mucoid strain (PAO1 *mucA22*; PDO300) and created a Δ*pqsA* allele in this background. In the macrocolony proximity assay, PDO300 induced STX production in the adjacent USA300 macrocolonies, although without growth inhibition (Figure 3C, left). However, for the PDO300Δ*pqsA* mutant, STX induction was abolished (Figure 3C, right). To quantify this, USA300 was grown planktonically in media supplemented with spent media from either PDO300, PDO300Δ*pqsA*, or PAO1Δ*pqsA*, followed by STX extraction and quantification. When grown in the presence of PDO300 spent media, USA300 produced significantly more STX, compared to growth in media alone (Figure 3D). Neither PAO1Δ*pqsA* nor PDO300Δ*pqsA* spent media was able to induce STX production. This suggests that mucoid *P. aeruginosa* also induces *S. aureus* STX production in an HQNO-dependent manner.

### *P. aeruginosa*-induced STX production protects *S. aureus* from H_2_O_2_-mediated killing

STX provides *S. aureus* resistance to oxidative stress, as a *crtM* mutant deficient in STX production was more sensitive to H_2_O_2_-mediated killing than WT *S. aureus*^20^. However, the survival of *S. aureus* to oxidative stress upon STX induction is not described. We, therefore, wanted to determine if *P. aeruginosa*-induced STX production could be beneficial for *S. aureus* survival in the presence of H_2_O_2_. As previously described, USA300 and *crtM*::Tn were grown overnight in media supplemented with either PAO1 or Δ*pqsA* spent media to induce STX production. The cultures were then subjected to 3% H_2_O_2_-mediated killing for up to 2h (Figure S4A, Figure 4), and *S. aureus* survival was quantified at designated time points compared to that of 0h. At 1h of H_2_O_2_ treatment, little difference in survival was observed for USA300 or *crtM*::Tn (Figure S4A). At 2h, the survival of H_2_O_2_-treated USA300 and *crtM*::Tn, grown without *P. aeruginosa* spent media, reduced to 49% and 29% respectively (Figure 4). Prior growth in media supplemented with PAO1 spent media, but not Δ*pqsA*, significantly increased USA300 survival by 4-fold, compared to USA300 grown in media alone. There was no significant difference in *crtM*::Tn survival under these conditions. The above data indicate that *P. aeruginosa*-induced STX production can further protect *S. aureus* from H_2_O_2_-mediated killing.

**Figure 4.**
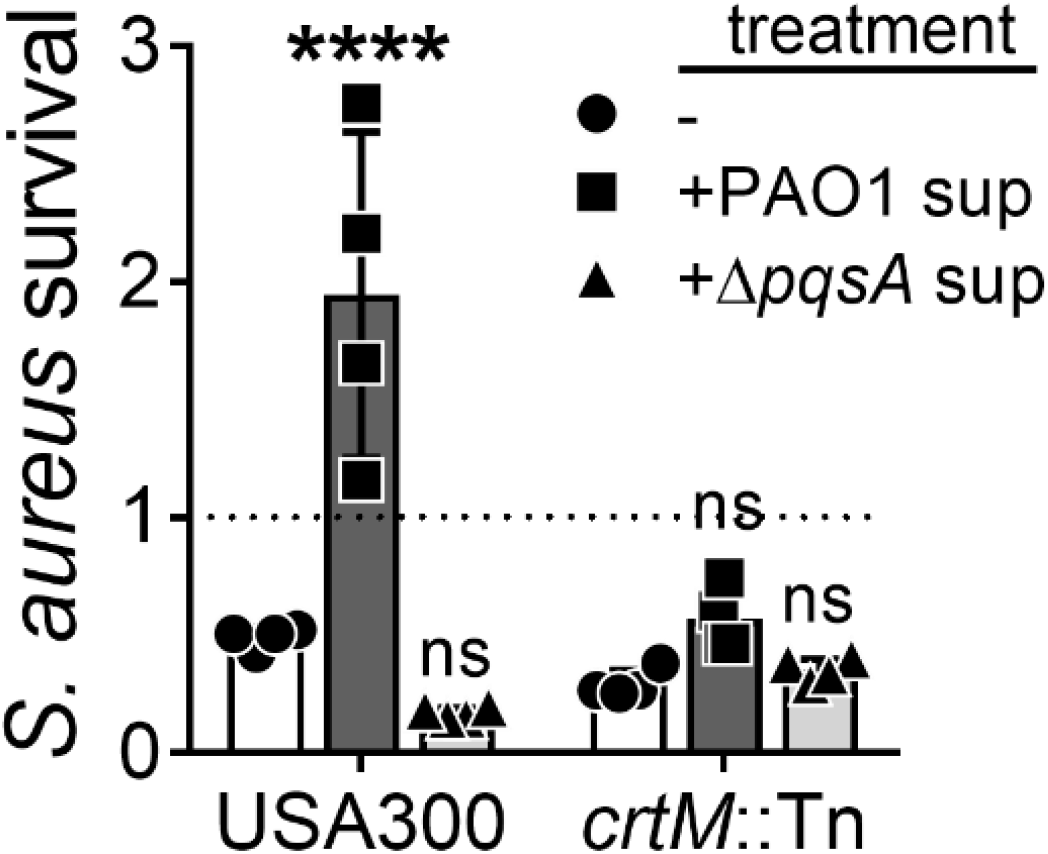
STX induction protects *S. aureus* from H_2_O_2_-mediated killing. *S. aureus* USA300 and *crtM::Tn* were pre-treated with or without 5% filter-sterilized PAO1 or Δ*pqsA* spent media (sup) overnight and then subjected to 3% H_2_O_2_-mediated killing for 2h. *S. aureus* survival is presented as CFUs normalized to the starting CFUs at 0h. Data are presented as mean ± SD from the results of at least 3 biological replicates, each with 3 technical replicates. ****, *P*<0.0001; ns, not significant, compared to the no treatment group (-).

### STX protects both mucoid and non-mucoid *P. aeruginosa* from H_2_O_2_-mediated killing

STX is a potent antioxidant and scavenges free radicals via conjugated double bonds^20,36^. We, therefore, hypothesized that in co-culture, STX could also protect *P. aeruginosa* from H_2_O_2_-mediated killing. PAO1 alone, or mixed with an equal amount of *S. aureus*, was treated with 3% H_2_O_2_ for 1h (Figure 5A). PAO1 survival increased >10-fold when mixed with USA300, compared to PAO1 alone. As expected, co-culture with *crtM*::Tn did not significantly change PAO1 survival in the presence of H_2_O_2_. The above data suggest that USA300 protects PAO1 from H_2_O_2_-mediated killing and this protection is STX-dependent.

**Figure 5.**
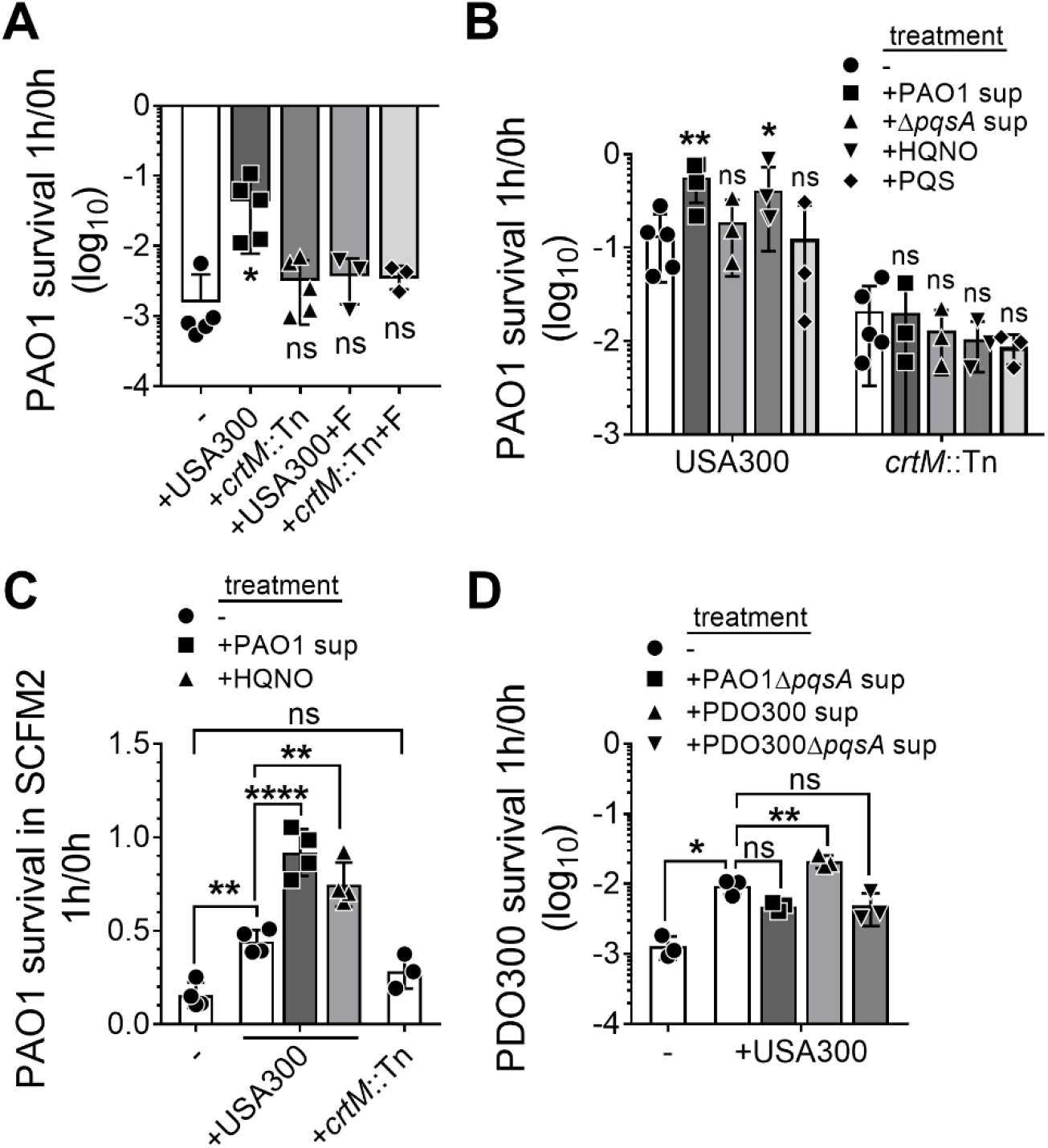
STX induction protects both mucoid and non-mucoid *P. aeruginosa* from H_2_O_2_-mediated killing. **(A)** PAO1, alone or mixed with an equal amount of *S. aureus*, was subjected to 3% H_2_O_2_-mediated killing for 1h. USA300 and *crtM*::Tn were pre-treated with 50μg/mL flavone (+F) to inhibit STX production and serve as controls. **(B, C)** PAO1, alone or mixed with an equal amount of *S. aureus* with various treatments, was subjected to 3% H_2_O_2_-mediated killing for 1h in either LB (B) or SCFM2 (C). USA300 and *crtM*::Tn were pre-treated with or without 5% filter-sterilized PAO1 or Δ*pqsA* spent media (sup), or 5μM HQNO or PQS overnight. **(D)** PDO300, either alone or mixed with an equal amount of USA300 with various treatments, was subjected to 3% H_2_O_2_-mediated killing for 1h. USA300 was pre-treated with or without 20% (v/v) filter-sterilized PAO1Δ*pqsA*, PDO300 or PDO300Δ*pqsA* spent media (sup). PAO1 and PDO300 survival is presented as CFUs normalized to the starting CFUs at 0h. Data are presented as mean ± SD from the results of at least 3 biological replicates, each with 3 technical replicates. *, *P* < 0.05; **, *P* < 0.01; ****, *P* < 0.0001; ns, not significant, compared to the *P. aeruginosa* alone group (A) or the no treatment group (B) unless stated in the figures.

We then wanted to confirm that the protection conferred to *P. aeruginosa* from H_2_O_2_-mediated killing by USA300 was due to the loss of STX production rather than other potential functions of *crtM*. To test this, USA300 and *crtM::Tn* were grown overnight in media supplemented with flavone, a plant flavonoid that inhibits *S. aureus* STX production without affecting growth^37^. The *S. aureus* cultures were then mixed with PAO1 and treated with 3% H_2_O_2_ for 1h, and bacterial survival was quantified (Figure 5A). Flavone completely abolished the protection that USA300 conferred on PAO1 against H_2_O_2_. No difference in the survival of PAO1 was observed when mixed with *crtM*::Tn grown with or without flavone. As expected with 1h treatment conditions, survival for both USA300 and *crtM*::Tn was not affected (Figure S4B), suggesting that the difference in PAO1 survival is attributed to STX, rather than the amount of *S. aureus* present. Together, the above data provide additional support that *S. aureus* STX enhances *P. aeruginosa* protection from H_2_O_2_-mediated killing.

Since HQNO induces *S. aureus* STX production, we further tested if *S. aureus* with induced STX production affords better protection to PAO1 from H_2_O_2_. USA300 STX was induced when grown in LB supplemented with PAO1 spent media or HQNO (Figure 1B, 2C). When mixed with the above cultures, the survival of PAO1 to H_2_O_2_-mediated killing was increased > 3-fold, compared to PAO1 mixed with USA300 grown in only LB (Figure 5B). Supplementing with Δ*pqsA* spent media or PQS during growth did not increase the ability of USA300 to protect PAO1. No significant difference was found in the survival of PAO1 when mixed with *crtM*::Tn grown with or without any *P. aeruginosa* exoproducts. The survival of USA300 and *crtM*::Tn remained unchanged for all conditions (Figure S4C). The above data indicate that induced STX production in *S. aureus* affords better protection to PAO1 from H_2_O_2_-mediated killing.

In addition to growth in LB, the presence of STX-producing USA300 also protected PAO1 from H_2_O_2_-mediated killing in SCFM2 (Figure 5C). Moreover, USA300 with increased STX production, induced by PAO1 spent media or HQNO, afforded better protection to PAO1 in SCFM2. The survival of USA300 remained unchanged for all conditions (Figure S4D). The above data suggest that *S. aureus* STX also protects *P. aeruginosa* from H_2_O_2_ killing in environments mimicking the CF lungs.

We also examined if STX can protect mucoid PDO300 from H_2_O_2_ killing (Figure 5D). Compared to PDO300 alone, the presence of USA300 increased the survival of PDO300 by 10-fold. PDO300 survival was further increased when the mixed USA300 was grown in LB supplemented with PDO300 spent media, in which case the STX production was induced (Figure 3D). This was not observed with PDO300Δ*pqsA* or PAO1Δ*pqsA* spent media, suggesting that USA300 with induced STX production affords better protection to PDO300 from H_2_O_2_-mediated killing. As expected, USA300 survival remained unchanged in all conditions (Figure S4E). The above data indicate that STX protection from H_2_O_2_-mediated killing extends to *P. aeruginosa* mucoid strains.

### STX protects *P. aeruginosa* from killing by human neutrophils

Neutrophils can combat pathogens by generating both intra- and extra-cellular reactive oxygen species (ROS)^38^. STX mediates *S. aureus* resistance to neutrophil killing by serving as an antioxidant^20,36^. In the experiments outlined above, we demonstrated that STX protected *P. aeruginosa* from oxidative stress generated by H_2_O_2_ (Figure 5). We wanted to determine if *S. aureus* STX could also protect *P. aeruginosa* from killing by human neutrophils. To test this, human peripheral blood-derived neutrophils were infected with fluorescently tagged USA300 and PAO1 for 1h, and bacterial survival was visualized by wide-field fluorescent microscopy. To inhibit STX production, USA300 was grown in the presence of flavone as described (Figure 5A). As shown in the representative images (Figure 6A) and quantifications of the fluorescent signal (Figure 6B), PAO1 survival was higher in the co-infection with USA300, but not flavone-treated USA300, compared to PAO1 mono-infection. PAO1 survival was also quantified by enumerating bacteria colony forming units (CFUs) before and after neutrophil exposure (Figure 6C). A significant increase was found in the survival of PAO1 when co-infected with USA300 (91%), but not with *crtM*::Tn (40%), compared to PAO1 alone (24%). *S. aureus* survival remained unchanged in mono-infections or co-infections with PAO1 (Figure S5). The above data suggest that *S. aureus* STX protects *P. aeruginosa* from killing by human neutrophils.

**Figure 6.**
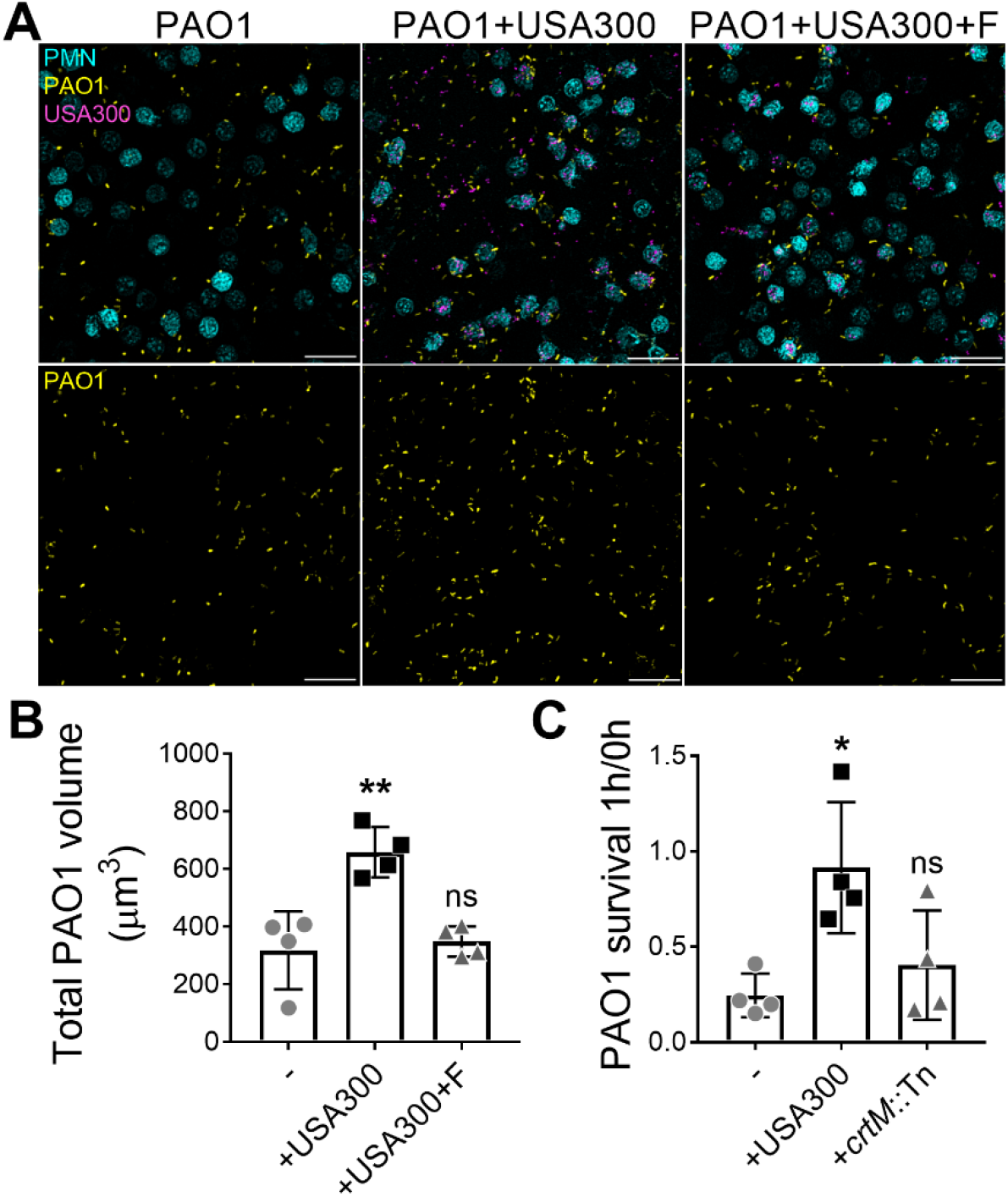
STX can protect *P. aeruginosa* from killing by human neutrophils. **(A, B)** PAO1-TdTomato, either alone or mixed with an equal amount of USA300-GFP, was subjected to adhered human neutrophil (PMN) for 1h to assess PAO1 survival (MOI = 10 for each species). USA300-GFP was pre-treated with 50μg/mL flavone (+F) to inhibit STX production. **(A)** Representative images of PAO1 and USA300 infected neutrophils. Scale bar: 20μm. **(B)** Total PAO1 volume was quantified by measuring fluorescence intensity. Data are presented as mean ± SD from the results of 4 biological replicates, each with 6 technical replicates. **(C)** PAO1, either alone or mixed with an equal amount of USA300 or *crtM*::Tn, was subjected to human neutrophil killing for 1h (MOI = 10 for each species). PAO1 survival is presented as CFUs normalized to the starting CFUs at 0h. Data are presented as mean ± SD from the results of 4 biological replicates, each with 3 technical replicates. *, *P* < 0.05; **, *P* < 0.01; ns, not significant.

### STX enhances *P. aeruginosa* infection *in vivo*

Our findings demonstrate that STX protects *P. aeruginosa* from killing by H_2_O_2_ and neutrophils (Figure 5,6). Since neutrophils are one of the first innate immune cells recruited to the infection site^39^, we hypothesized that STX could protect PAO1 during the early stages of infection *in vivo*. To test this, a dermal full-thickness murine wound model was used to examine *P. aeruginosa* and *S. aureus* co-infection^40^. Briefly, two dorsal wounds were generated by punch biopsies and infected with either a luminescent tagged PAO1 strain, USA300, *crtM*::Tn, or both species. The infection was allowed to progress for 3 days to focus on early bacterial infection and innate immunity (Figure 7A). By using an *in vivo* imaging system (IVIS), we were able to monitor the burden of the bioluminescent PAO1 daily by measuring signal intensity. PAO1 burden remained the highest in mice co-infected with PAO1 and USA300 throughout the 3-day infection period, compared with PAO1 mono-infection and *crtM*::Tn co-infection (Figure 7BD). By quantifying the area under the curve (AUC) of Figure 7D, significant increases were found when comparing USA300 co-infection to PAO1 mono-infection and *crtM*::Tn co-infection. No difference was found between PAO1 mono-infection and *crtM*::Tn co-infection (Figure S6A). The above data suggest that *S. aureus* STX increases the PAO1 burden throughout infection.

**Figure 7.**
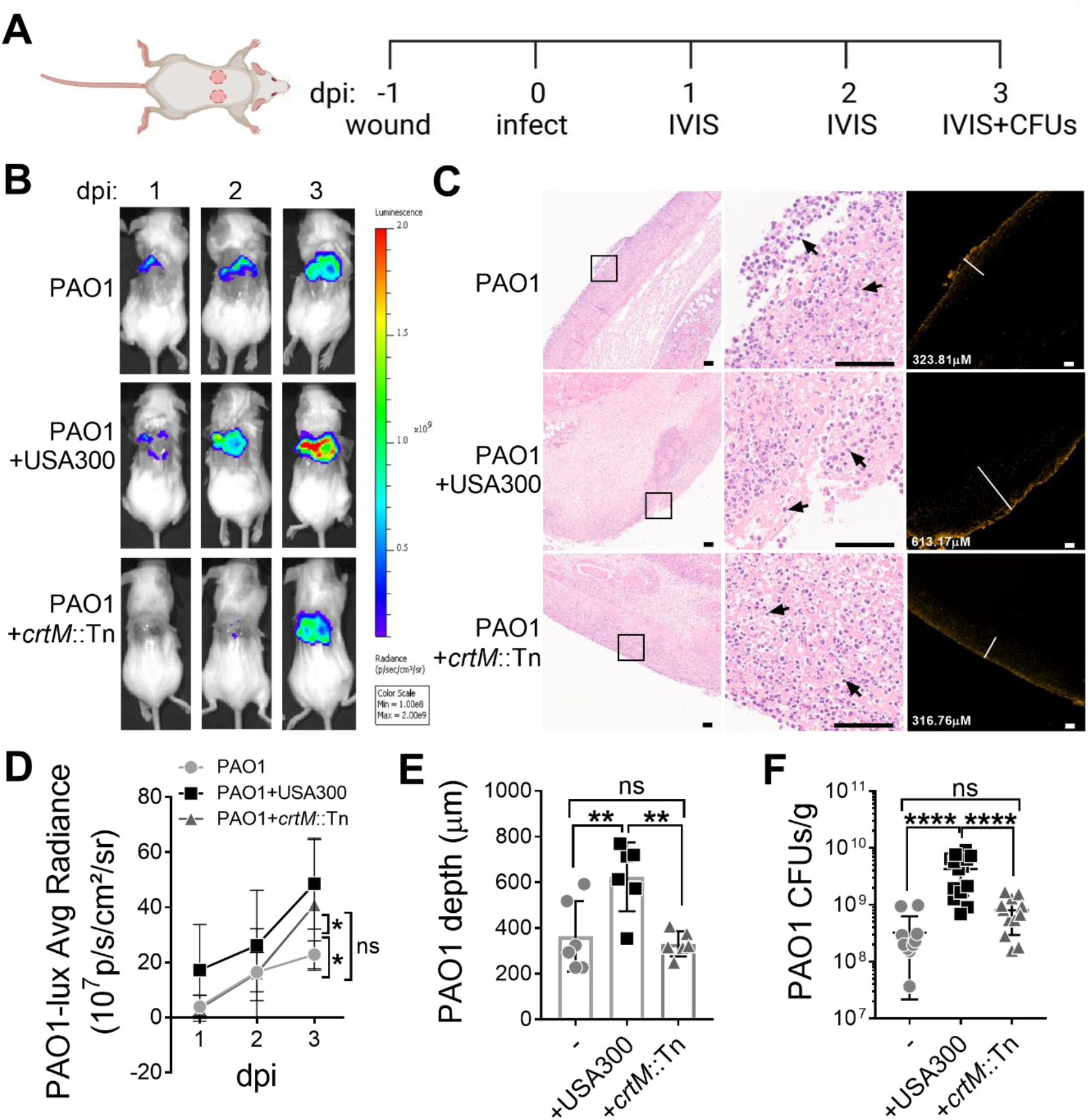
STX promotes the establishment of *P. aeruginosa* infection *in vivo*. **(A)** Schematic of the murine wound model and course of infection. 2 identical full-thickness dorsal wounds were generated using 6mm punch biopsies. After 24h, the wounds were mono-infected with luminescent PAO1, USA300, or *crtM*::Tn, or co-infected with PAO1 and USA300 or *crtM*::Tn. On 1, 2, and 3 days post-infection (dpi), IVIS was used to monitor PAO1 burden among all groups. Wounds were harvested and homogenized to plate for both PAO1 and *S. aureus* CFUs on 3 dpi. **(B)** Representative images of luminescent PAO1 detected using IVIS on murine wounds throughout the 3 days of infection. **(C)** Representative images of H&E (left and middle panels) and IF (right panel) stained adjacent wound sections (4μm). The left panel shows the wound beds in low magnification. Magnified boxed areas are shown in the middle panel, with black arrows pointing to neutrophil infiltration. The right panel shows the presence of immunofluorescently-labeled PAO1 and the white lines measure the depth of PAO1 penetration (labeled in the bottom left corner). Scale bar: 100μm. **(D)** Luminescent signal intensity of PAO1-lux was quantified by the average radiance. Significant differences were determined by comparing the area under the curve (Figure S6A). Data presented as mean ± SD from the results of more than 12 biological replicates. **(E)** The depth of PAO1 penetration into the wound with PAO1 mono-infection or co-infection with USA300 or *crtM*::Tn. Data are presented as mean ± SD from the results of 6 biological replicates. **(F)** PAO1 CFUs per gram of wound tissue (CFUs/g) among all groups was quantified. Data are presented as mean ± SD from the results of more than 12 biological replicates with 3 technical replicates. *, *P* < 0.05; **, *P* < 0.01; ****, *P* < 0.0001; ns, not significant.

On day 3 post infection, wound tissues were harvested and processed for either hematoxylin and eosin (H&E) staining or immunofluorescence (IF) labeling of PAO1, to assess inflammation and PAO1 localization respectively. We observed infiltration of mixed inflammatory cells, which predominantly consisted of neutrophils, into the wound bed for all groups (Figure 7C, left and middle panel), close to the location of PAO1 (Figure 7C, right panel). IF staining showed that PAO1, when co-infected with USA300, penetrated significantly deeper into the tissue (623.8 ± 150.9μm) compared to mono-infection (363.5 ± 154.8μm) or co-infection with *crtM*::Tn (330.2 ± 54.81μm) (Figure 7CE). Total PAO1 fluorescent signal from immunofluorescent images was quantified as another indication of PAO1 burden and was higher when PAO1 was co-infected with USA300, but not *crtM*::Tn or PAO1 alone (Figure S6B). The above results were corroborated by enumerating bacteria after 3 days of infection (Figure 7F). When PAO1 was co-infected with USA300 but not *crtM*::Tn, there was a significant 10-fold increase in PAO1 burden, compared to PAO1 mono-infection. Overall, the above data indicate that STX-producing *S. aureus* promotes *P. aeruginosa* wound colonization during the early stages of infection, implying a role for STX in enhancing *P. aeruginosa* infection *in vivo*.

## DISCUSSION

In this study, we identified a novel role for *P. aeruginosa* HQNO in inducing *S. aureus* STX production (Figure 1-3). The well-studied influence of HQNO on *S. aureus* is that, at a much higher concentration (400μM), it antagonizes growth by inhibiting respiration^27,41,42^, and promotes the formation of *S. aureus* small colony variants^12,27,42^, which are slow-growing and non-pigmented^43^. However, we observed increased STX production in the presence of sub-lethal HQNO (5μM) concentrations. It is likely that varying concentrations of HQNO may elicit different responses in *S. aureus*. Many factors contribute to the local concentration of HQNO. The spatial distribution of bacteria can alter local QS signal concentration^46^. In Figure 1A, USA300 growth was inhibited by the adjacent PAO1 with high local HQNO concentration (black arrow). The antagonism disappeared and STX induction was observed as the HQNO concentration decreased, due to the increased distance between the two bacteria macrocolonies (orange arrows). As the distance between the two colonies further increased, HQNO concentration was likely too low to induce STX production (yellow arrows). In addition, different *P. aeruginosa* laboratory and clinical strains can produce varying amounts of HQNO, ranging from 0 to 50μM^32,33^. Mucoid *P. aeruginosa*, with an overproduction of alginate, is commonly isolated from chronic infections and can co-exist with *S. aureus* better than the non-mucoid counterparts^35^. Exogenous alginate can mitigate the killing of *S. aureus* by down-regulating HQNO production and other antagonistic factors in both mucoid and non-mucoid *P. aeruginosa*^44^. Consistent with this, we observed reduced USA300 antagonism and induction of STX in some of the mucoid clinical isolates and PDO300 in an HQNO-dependent manner (Figure 3BCD). The above evidence suggests that mucoid *P. aeruginosa* can create a low-HQNO environment that promotes *S. aureus* niche compatibility. This allows for the induction of STX which is beneficial for both species. In addition to mucoid strains, some non-mucoid *P. aeruginosa* clinical isolates can also induce STX production without inhibiting the growth of adjacent USA300 (Figure 3B), possibly due to reduced HQNO production in these isolates. Apart from spatial distribution and strain variations, nutrient availability and host factors may also alter HQNO concentration^45-47^. Overall, we predict that different concentrations of *P. aeruginosa* HQNO may be responsible for controlling the balance between cooperative and competitive behaviors between *P. aeruginosa* and *S. aureus*.

The mechanism(s) of how HQNO induces STX remains unclear. HQNO can activate the expression of the *S. aureus* global transcriptional regulator sigma factor B (SigB)^27^, which regulates STX production ^48^. In addition, the induction seems to correlate with but is not dependent on growth inhibition, as most of the STX-inducing *P. aeruginosa* clinical isolates can inhibit *S. aureus* growth (Figure 3B). This implies that STX may be induced in response to HQNO-mediated antagonism. HQNO can interfere with the electron transfer system in bacteria and mitochondria, which results in a leak of reducing equivalents to O_2_ and the production of ROS^49-51^. Although not demonstrated for *S. aureus*, an excessive amount of HQNO can result in ROS production and be self-poisoning for *P. aeruginosa*^52^. It is possible that *S. aureus* produces more STX to counteract HQNO-generated ROS. However, results from our experiments suggest otherwise. PAO1-mediated growth inhibition of USA300 and *crtM*::Tn were comparable (Figure 1A, black arrows). When co-cultured planktonically overnight with PAO1, no significant difference was observed between USA300 and *crtM*::Tn survival (Figure S7). Moreover, no STX induction was found in USA300 when treated with ciprofloxacin, even though it also results in ROS production ^53^ (Figure S1A). We speculate that the induction of STX by HQNO may be a result of *S. aureus* sensing HQNO signals specifically. Interspecies signaling is a key factor contributing to polymicrobial interaction and spatial distribution. There is evidence that *S. aureus* may have membrane receptors for another *P. aeruginosa* QS molecule, acyl homoserine lactone^54,55^, but little is known about HQNO. Interestingly, STX is produced by *S. aureus* class III clinical isolates but is not induced by the adjacent PAO1 (Figure 3A). This suggests that these isolates may be blind to sensing HQNO. By examining the clinical isolates’ genomes and using STX induction as an output to screen for HQNO-unresponsive variants, future studies will uncover the mechanism(s) of how *S. aureus* senses *P. aeruginosa* HQNO and elevates STX production.

STX is an antioxidant and scavenges free radicals^36^. In accord with this, it not only protected *S. aureus* but also *P. aeruginosa* from H_2_O_2_-dependent killing (Figure 4, 5). The protection was also apparent when bacteria were subjected to neutrophil killing (Figure 6). Other than ROS, neutrophils can release APs to eliminate bacterial infections^56^. STX is positively associated with *S. aureus* cell membrane fluidity and resistance to neutrophil-derived APs^21^. However, this is unlikely how STX protects PAO1 from neutrophil killing, as it would make the APs more available to PAO1, resulting in decreased PAO1 survival. Apart from the *in vitro* and *ex vivo* experiments, we also examined *P. aeruginosa* and *S. aureus* co-infection *in vivo*. Co-infection correlates with increased disease severity, compared to mono-infections of either species^4-6^. Co-infection of murine wounds with STX-producing USA300 promoted PAO1 infection much greater than the STX-deficient *crtM*::Tn (Figure 7). Interestingly, despite the increased burden of PAO1, little difference was observed in the disease state of the wounds, determined by pathology scoring (Figure S6C). Given the results from *in-vitro* and *ex-vivo* experiments, we speculate that the *in-vivo* fitness benefit afforded to PAO1 by USA300 is due to the antioxidant nature of STX. However, we cannot exclude the possibility that STX is not the only factor produced by USA300 that enhances PAO1 infection.

Apart from PAO1, we also quantified the survival of USA300 and *crtM*::Tn in the presence of ROS. Liu *et al* found that a *crt* mutant had decreased survival compared to WT *S. aureus* when subjected to H_2_O_2_ and neutrophil killing, as well as lower fitness in a subcutaneous abscess murine model^20^. In accord with this, *crtM*::Tn survival, compared to USA300, was modestly lower when subjected to H_2_O_2_ killing for 2h (Figure 4) and in the murine wound infection (Figure S6D). However, no difference was found between the survival of the two strains when infecting neutrophils (Figure S5). These discrepancies are likely due to different experimental conditions used in these studies^20^.

Since HQNO can induce STX production which is beneficial for USA300 to resist H_2_O_2_ killing (Figure 1, 2, 4), we also examined the reciprocal: if the presence of PAO1 was beneficial for USA300 *in vivo. S. aureus* survival during co-infections with *P. aeruginosa* can be affected by many factors. Host immunity and antagonism from *P. aeruginosa* can reduce *S. aureus* survival, while potential STX induction from *P. aeruginosa* can be beneficial for *S. aureus*. We found that the survival of both USA300 and *crtM*::Tn was reduced during co-infection with PAO1, compared to mono-infections (Figure S6D). This suggests that the presence of *P. aeruginosa*, overall, is detrimental to *S. aureus* survival *in vivo*. However, the *S. aureus* burden remained high (10^7^∼10^8^ CFUs/g) after 3 days of co-infection with PAO1. This is consistent with previous findings that the *in-vivo* wound environment can promote co-existence despite their antagonistic relationship *in vitro*^3,47,57^. Interestingly, the ratio of *crtM*::Tn CFUs to that of USA300, in the co-infections with PAO1, was lower compared to *S. aureus* mono-infections (Figure S6E). Given that *crtM*::Tn was not more sensitive to antagonism by PAO1 than USA300 (Figure S7), we speculate that this difference may be attributed to the relatively higher survival of USA300 than *crtM*::Tn, when co-infected with PAO1. The presence of PAO1 may induce STX production in USA300 which promotes its resistance to host ROS resulting in better survival, but not in *crtM*::Tn. Unfortunately, we were unable to extract and directly quantify STX levels from the homogenized wounds due to contamination of host debris. Overall, our finding adds another layer to the already complex interaction between *P. aeruginosa* and *S. aureus in vivo*.

In summary, we identified a novel role for *P. aeruginosa* HQNO in inducing *S. aureus* STX production which is prevalent among clinical isolates. We also discovered a new cooperative behavior between the two pathogens during co-infections, resulting in resistance to host ROS and increased *P. aeruginosa* burden *in vivo*. Our study contributes to the understanding of complex polymicrobial interactions and highlights the need for future investigations in treating polymicrobial infections.

## MATERIALS AND METHODS

### Bacterial strains and growth conditions

All bacterial strains and plasmids are listed in Table S1. All planktonic cultures were grown at 37°C with 200rpm shaking for 16h. *S. aureus* planktonic culture was grown in either lysogeny broth (10g/L tryptone, 5g/L yeast extract, 10g/L NaCl; LB) or SCFM2^58^. *P. aeruginosa* planktonic culture was grown in either LB with no salt (LBNS) or SCFM2. For macrocolony proximity assay,*P. aeruginosa* and *S. aureus* were grown on lysogeny agar (LB supplemented with 1.5% agar; LA).

### Macrocolony proximity assay

Macrocolonies were grown by inoculating 5µl of overnight bacterial culture onto the surface of LA in two sets of experiments. 1) *P. aeruginosa* and *S. aureus* were spotted on opposite sides of the plate with distances of 0cm, 1cm, 2cm, and 3cm in between. 2) 5µl of either PAO1 culture, 50µM HQNO or PQS, the solvent (methanol and ethanol), or antibiotics (10mg/mL daptomycin or 1mg/mL ciprofloxacin) was spotted onto the center of the plate, and the USA300 macrocolonies were spotted 0.5cm, 1cm, 2cm and 3cm away from the center. The plate was incubated at 37°C overnight and the *S. aureus* colonies were examined for their survival and pigment production.

### STX production in planktonic culture

*P. aeruginosa* overnight cultures were normalized to OD_600_ 2.5, and filter sterilized to collect cell-free spent media. To induce STX production, early-stationary phase *S. aureus* cultures (OD_600_ = 1) were supplemented with either 5% or 20% (v/v) *P. aeruginosa* spent media or 5µM of PQS or HQNO (Cayman Chemicals). For PAO1 and its variants, 5% spent media (v/v) was added to *S. aureus*. Since PDO300 produces less HQNO^35^, 20% spent media (v/v) of PDO300 and its variants was added to *S. aureus*. To inhibit STX production, *S. aureus* was grown in LB supplemented with 50μg/mL flavone (Sigma-Aldrich) overnight.

### STX extraction

Extraction and quantification of STX were carried out as previously described^17^ with modifications. In brief, overnight cultures of *S. aureus* were normalized to an OD_600_ of 3. After centrifugation, the pellet was resuspended in 250µl of methanol and incubated at 55°C for 3min. The samples were centrifuged to collect the supernatant and measured for absorbance at OD_440_, OD_462_, and OD_491_ using a plate reader (SpectraMax® i3x; Molecular Device).

### H_2_O_2_-mediated killing assay

This assay was carried out as previously described^20^ with modifications. Overnight cultures of *P. aeruginosa* and *S. aureus* were diluted to OD_600_ 0.5 in fresh LB. They were either combined at a 1 : 1 ratio or separately subjected to 3% H_2_O_2_ (Spectrum Chemical) and incubated at 37°C with 200rpm shaking for up to 2h. Aliquots were taken at every hour, treated with 2000U/mL catalase (Sigma-Aldrich), serially diluted, and plated on Difco™ Pseudomonas Isolation Agar (PIA) and BBL™ Mannitol Salt Agar (MSA) to enumerate for CFUs of *P. aeruginosa* and *S. aureus*, respectively. Bacterial survival at each time point was normalized to the CFUs at 0h.

### Neutrophil isolation

Informed written consent was obtained from all 4 healthy donors before the collection of peripheral blood for isolating primary human neutrophils. All procedures were approved by the Ohio State University Institutional Review Board (IRB-2009H0314). Neutrophils were isolated as previously described^59^. Briefly, heparinized blood from healthy human donors was collected in saline. Ficoll-Paque® PLUS (GR Healthcare) was layered on top of the blood and then centrifuged at 404 × *g* for 40min at 23°C. The pellet was then resuspended in an equal volume of 3% cold Dextran in 0.9% NaCl and allowed for sediment for 20min on ice. The upper layer was centrifuged at 665 × *g* for 10min at 4°C. The resulting pellet was resuspended in cold endotoxin-free H_2_O for 30s to lyse red blood cells before 1.8% NaCl solution was immediately added to restore isotonicity. The sample was centrifuged at 131 × *g* for 3min at 4°C, and the pellet containing neutrophils was resuspended in HBSS (without calcium, magnesium, or phenol red; Corning) and counted in a hemocytometer chamber^60^.

### Neutrophil killing assay

This assay was carried out as previously described ^61^ with modifications. *P. aeruginosa* and *S. aureus* overnight cultures were normalized to an OD_600_ of 0.5, washed with HBSS, and opsonized with 20% human serum (CompTech) for 30min at 37°C. The two bacteria were then either mixed at a 1 : 1 ratio or separately incubated with neutrophils statically for 1h at 37°C (MOI = 10 for each bacterial species). The samples were centrifuged at 18000 x *g* for 10min to lyse the neutrophils and release internalized bacteria. The pellets were resuspended in HBSS, serially diluted, and plated on PIA and MSA to enumerate CFUs. Bacterial survival was normalized to the CFUs at 0h.

For microscopy analysis, neutrophils (2 × 10^6^ cells per well) were seeded on poly-l-lysine coated coverslips in HBSS supplemented with 100 μM CellTracker™ Blue (Invitrogen) for 30min at 37°C, 5% CO_2_. Fluorescently tagged *P. aeruginosa* (PAO1-TdTomato)^62^ was grown overnight in LB supplemented with 300ug/mL of carbapenem. Fluorescently tagged *S. aureus* (USA300-GFP)^63^ was grown overnight in LB supplemented with or without 50μg/mL flavone. Attached neutrophils were infected with PAO1-TdTomato, USA300-GFP, or both species for 1h at 37°C, 5% CO_2_ (MOI = 10 for each bacterial species). Unattached cells were washed away with HBSS. Coverslips were fixed in 4% paraformaldehyde for 30min at room temperature, mounted to slides using Prolong™ Gold antifade reagent (Invitrogen), and visualized using a Nikon Ti2 wide field microscope fitted with a 60x oil objective. 6 images with Z-stack were taken for each sample in every experiment. By using the NIS-elements AR software, the images were clarified, deconvoluted, and thresholded to quantify the total volume of bacteria. Representative images shown in Figure 6C were focused 2D images, created from the original 3D images by the software.

### Dermal full-thickness murine wound infection

This assay was carried out as previously described^40^ with modifications. All procedures were approved by the Ohio State University Institutional Animal Care and Use Committee (2017A00000028-R1; 2008R0135-R1; 2011R00000021-R1). Briefly, 6-week-old female BALB/c mice were anesthetized using isoflurane gas, and the dorsal area was shaved. The dorsal area was then sterilized with ethanol and two identical full-thickness dorsal wounds were generated with a 6mm punch biopsy tool (Integra™ Miltex®) and bandaged with a Tegaderm dressing (3M). The mice were allowed to recover for 24h before infection. For infection, mid-log *P. aeruginosa* and *S. aureus* (OD_600_ = 0.5) were washed and resuspended in 0.9% endotoxin-free saline. Each wound was infected with bacterial cultures containing 5×10^6^ cells of either PAO1 (containing a constitutively expressed luminescent marker^64^), USA300 or *crtM*::Tn, or both species. A total of 7 animals were used for each group. To assess PAO1 burden throughout infection, mice were anesthetized, and the wound luminescence was imaged daily with an IVIS Lumina II optical imaging system (PerkinElmer Inc.). The acquired images were scaled to the radiance of 1e8 to 2e9. The average radiance of PAO1-lux on each animal was used to access the PAO1 burden throughout infection. Three days post infection, mice were euthanized by CO_2_ inhalation. The wounded tissue was collected, weighed, and placed in separate tubes containing 1 mL of PBS. All samples were homogenized with a Pro Scientific Bio-Gen Series Pro200 hand-held homogenizer for 45 s. The resulting solutions were serially diluted, plated on PIA and MSA, and incubated at 37 °C overnight. CFUs were calculated per gram of tissue.

### H&E and IF staining, and pathology analysis on the wound tissues

3 days post infection, wounds were harvested, fixed in 4% paraformaldehyde for a week, transferred into 100% ethanol, and sent to HistoWiz. The tissues were embedded in paraffin, sectioned longitudinally (4μm), and stained with H&E. Digital skin sections were subjectively assessed by HistoWiz for the severity and extent of inflammation to provide pathology scoring.

As for the IF staining, the slides were deparaffinized and blocked with 3% bovine serum albumin supplemented with 50 mM glycine, 0.05 % Tween20, and 0.1% Triton X-100 at 4°C overnight. Slides were then incubated with primary *P. aeruginosa* antibody^65^ (1 : 500 dilution) at 4°C overnight and secondary antibody (Alexa Fluor™ 647 chicken anti-rabbit IgG, Invitrogen; 1 : 500 dilution) at room temperature for 1h. They were visualized by microscopy (Nikon ECLIPSE Ti2) using a 4x objective. 6 wounds were imaged for each group. The depth of PAO1 penetration into the wound and total pixel count were measured by NIS-elements AR software.

### *P. aeruginosa* and *S. aureus* planktonic co-culture

Overnight cultures of PAO1 and USA300 or *crtM*::Tn were diluted to OD_600_ 0.05 and combined at a ratio of 1:1 in LB. The co-culture was incubated at 37°C shaking at 200rpm for 24h. It was serially diluted and plated on MSA to enumerate for *S. aureus* CFUs. *S. aureus* survival was normalized to the CFUs at 0h.

### Statistical analysis

Statistical significance was determined using either a One-way or Two-way ANOVA with Tukey’s post-hoc test. Analyses were performed using GraphPad Prism v.7 (Graphpad Software). Statistical significance was determined using a *P*-value <0.05.

## ACKNOWLEDGEMENTS

This study was supported by the National Institute of Health R01AI143916 to DJW and YL, R01AI077628 to PSR, R01AI134895 to EAM, and a fellowship program for Advancing Research in Infection and Immunity to YL. This work was supported in part by the Cure CF Columbus Translational Core (C3TC). C3TC is supported by the Division of Pediatric Pulmonary Medicine, the Biopathology Center Core, and the Data Collaboration Team at Nationwide Children’s Hospital. Grant support was provided by The Ohio State University Center for Clinical and Translational Science (National Center for Advancing Translational Sciences, Grant UL1TR002733) and by the Cystic Fibrosis Foundation (Research Development Program, Grant MCCOY19RO). We thank Dr. Traci Wilgus for helping us with the paraffin removal of wound sections.

