## Supplemental data for "Cross-Species Protection to Innate Immunity Mediated by A Bacterial Pigment"

**Table S1 Strains**

| Strains | Description | Source |
|---|---|---|
| *P. aeruginosa* laboratory strains | | |
| PAO1 | WT *P. aeruginosa* | [1] |
| Δ*pqsA* | PAO1 *pqsA* deletion mutant | [2] |
| PDO300 | Mucoid, a *mucA* derivative of PAO1 | [3] |
| PDO300Δ*pqsA* | Mucoid, *pqsA* deleted in PDO300 | this study |
| PAO1-TdTomato | PAO1 carrying a constitutively expressed Td-tomato-producing plasmid | [4] |
| PAO1-lux | Luminescent PAO1 | [5] |
| *Δ*P*psl* | PAO1 *psl* production deficient; *psl* operon promoter deletion mutant | [1] |
| Δ*pvdA* | PAO1 *pvdA* deletion mutant | [2] |
| *rhlA*::Tn | *rhlA* transposon mutant (UWGC:PW6886, PA3479::IS*phoA*/hah) | [6] |
| *lasA*::Tn | *lasA* transposon mutant (UWGC:PW4282, PA1871::IS*lacZ*/hah) | [6] |
| *P. aeruginosa* clinical isolates | | |
| 6546 | CF clinical isolate, mucoid | this study |
| 6547 | CF clinical isolate | this study |
| 6548 | CF clinical isolate | this study |
| 6550 | CF clinical isolate, mucoid | this study |
| 6551 | CF clinical isolate, mucoid | this study |
| 6559 | CF clinical isolate | this study |
| 6560 | 0CH5M4, CF clinical isolate | [2] |
| 6561 | 0CH7HJ, CF clinical isolate | [2] |
| 6565 | 0CHBKC, CF clinical isolate | [2] |
| 6566 | 0CHBKD, CF clinical isolate | [2] |
| 6354 | Wound isolate | this study |
| 6355 | Wound isolate | this study |
| 6356 | Wound isolate | this study |
| 6357 | Wound isolate | this study |
| 6358 | Wound isolate | this study |
| 6359 | Wound isolate | this study |
| 6360 | Wound isolate | this study |
| 6361 | Wound isolate | this study |
| 6362 | Wound isolate | this study |
| 6363 | Wound isolate | this study |
| 6364 | Wound isolate | this study |
| 6365 | Wound isolate | this study |
| 6366 | Wound isolate | this study |
| 6367 | Wound isolate | this study |
| 2901 | CF clinical isolate | this study |

| | | |
|---|---|---|
| 2902 | CF clinical isolate, mucoid | this study |
| 2903 | CF clinical isolate | this study |
| 2905 | CF clinical isolate, mucoid | this study |
| 2906 | CF clinical isolate | this study |
| *S. aureus* laboratory strains | | |
| USA300 | WT *S. aureus* | [7] |
| MSSA | ATCC 29213, Methicillin sensitive *S. aureus* | ATCC |
| USA300-GFP | USA300 with constitutively expressed GFP on the chromosome | [8] |
| *crtM*::Tn | *crtM* transposon mutant (NE1444, NARSA) | [7] |
| *S. aureus* clinical isolates | | |
| 6538 | CF clinical isolate | this study |
| 6539 | CF clinical isolate | this study |
| 6540 | CF clinical isolate | this study |
| 6541 | CF clinical isolate | this study |
| 6542 | CF clinical isolate | this study |
| 6543 | CF clinical isolate | this study |
| 6544 | CF clinical isolate | this study |
| 6545 | CF clinical isolate | this study |
| 6553 | CF clinical isolate | this study |
| 6554 | CF clinical isolate | this study |
| 6555 | CF clinical isolate | this study |
| 6556 | CF clinical isolate | this study |
| 6557 | CF clinical isolate | this study |
| 6558 | CF clinical isolate | this study |
| 6562 | CF clinical isolate | this study |
| 6563 | CF clinical isolate | this study |
| 6564 | CF clinical isolate | this study |
| 6567 | CF clinical isolate | this study |
| 6569 | CF clinical isolate | this study |
| 6585 | CF clinical isolate | this study |
| 6586 | CF clinical isolate | this study |
| 6587 | CF clinical isolate | this study |
| 6588 | CF clinical isolate | this study |
| 6589 | CF clinical isolate | this study |
| 6590 | CF clinical isolate | this study |
| 6591 | CF clinical isolate | this study |
| 6592 | CF clinical isolate | this study |
| 6593 | CF clinical isolate | this study |
| 6594 | CF clinical isolate | this study |
| 6595 | CF clinical isolate | this study |
| 6596 | CF clinical isolate | this study |

| | | |
|---|---|---|
| 6637 | CF clinical isolate | this study |
| 4101 | Bloodstream isolate | this study |
| 4102 | Bloodstream isolate | this study |
| 4103 | Bloodstream isolate | this study |
| 4104 | Bloodstream isolate | this study |
| 4105 | Bloodstream isolate | this study |
| 4106 | Bloodstream isolate | this study |
| 4107 | Bloodstream isolate | this study |
| 4108 | Bloodstream isolate | this study |
| 4109 | Bloodstream isolate | this study |
| 4110 | Bloodstream isolate | this study |
| 4111 | Bloodstream isolate | this study |
| 4112 | Bloodstream isolate | this study |
| 4113 | Bloodstream isolate | this study |
| 4114 | Bloodstream isolate | this study |
| 4115 | Bloodstream isolate | this study |
| 4116 | Bloodstream isolate | this study |
| 4117 | Bloodstream isolate | this study |
| 4118 | Bloodstream isolate | this study |
| 4119 | Bloodstream isolate | this study |
| 4120 | Bloodstream isolate | this study |
| 4121 | Bloodstream isolate | this study |
| 4122 | Bloodstream isolate | this study |
| 4123 | Bloodstream isolate | this study |
| 4124 | Bloodstream isolate | this study |
| 4125 | Bloodstream isolate | this study |
| 4126 | Bloodstream isolate | this study |
| 4127 | Bloodstream isolate | this study |
| 4128 | Bloodstream isolate | this study |
| 4129 | Bloodstream isolate | this study |
| 4130 | Bloodstream isolate | this study |

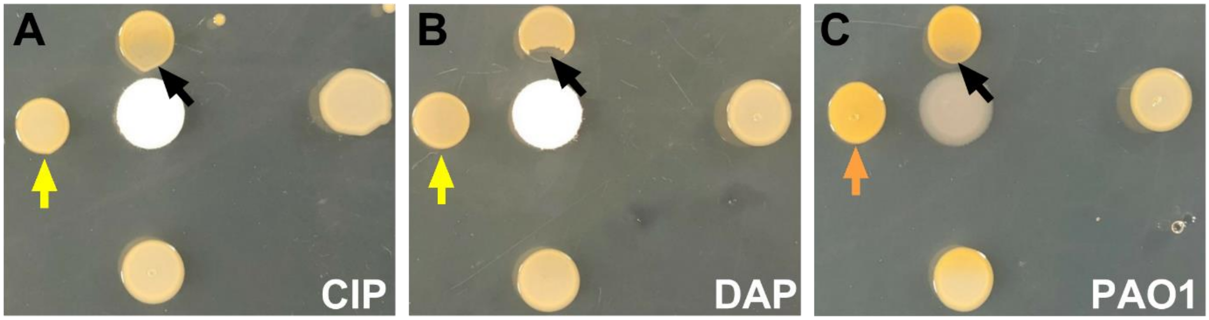


**Figure S1. *S. aureus* STX production is not induced by antibiotic-mediated growth inhibition.** USA300 was grown at different distances to discs soaked in 5μL of 1mg/mL ciprofloxacin (CIP), 10mg/mL daptomycin (DAP) or PAO1 overnight culture on solidified media in a macrocolony proximity assay. Yellow arrows point to USA300 colonies with no color change, orange arrows point to USA300 colonies with increased yellow pigmentation and black arrows point to USA300 colonies with inhibited growth.

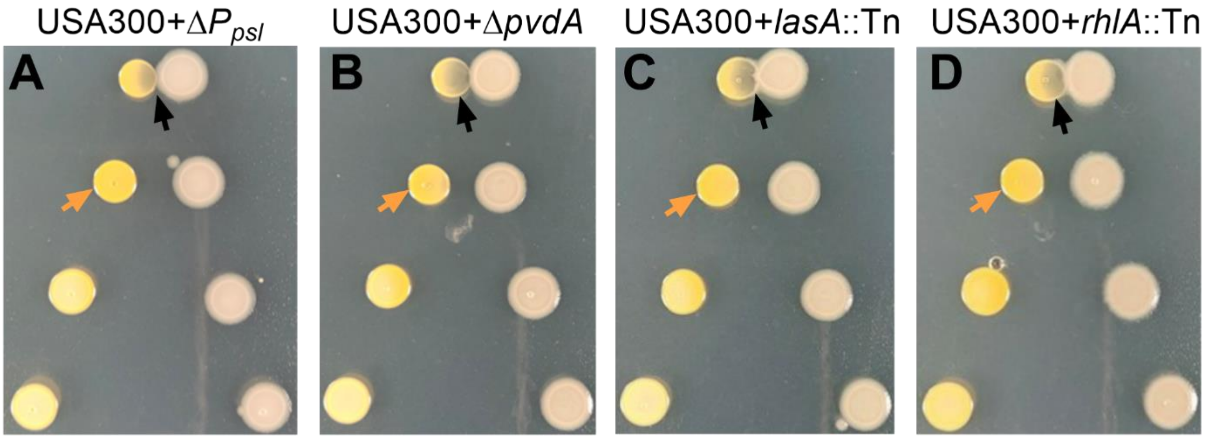


**Figure S2. *S. aureus* STX production when grown with PAO1 mutants with decreased antagonism towards *S. aureus*.** USA300 was grown at different distances to PAO1 mutants deficient in producing exopolysaccharide Psl ($\Delta P_{psl}$), pyoverdine (Δ*pvdA*), protease LasA (*lasA*::Tn) or rhamnolipid (*rhlA*::Tn) on solidified media in a macrocolony proximity assay. The orange arrows point to USA300 with increased yellow pigmentation, and the black arrows point to USA300 growth inhibition by PAO1.

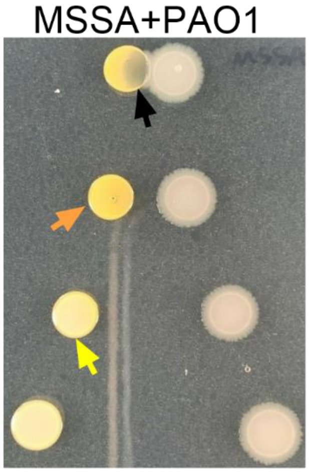


**Figure S3. Methicillin-sensitive *S. aureus* STX production is induced by *P. aeruginosa*.** MSSA was grown at different distances to PAO1 on solidified media in a macrocolony proximity assay. The orange arrow points to MSSA with increased yellow pigmentation, yellow arrow points to MSSA with no color change, and the black arrow points to MSSA growth inhibition by PAO1 .

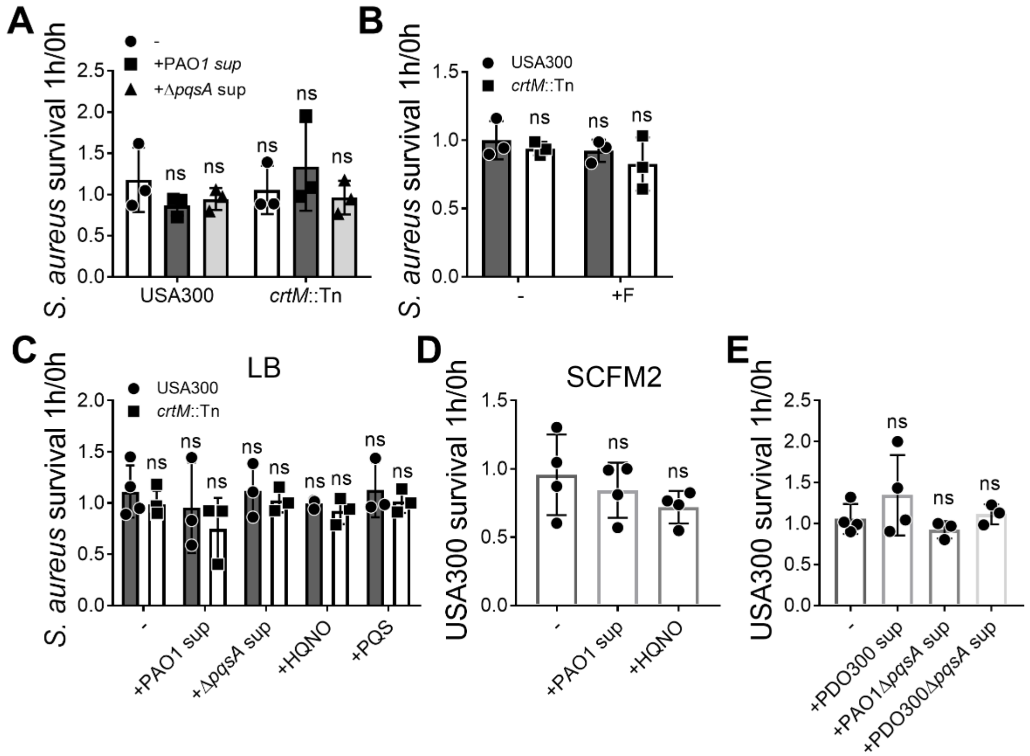


**Figure S4. *S. aureus* survial when treated with $H_2O_2$. (A)** USA300 and *crtM*::Tn were pre-treated with or without 5% filter-sterilized PAO1 or PAO1Δ*pqsA* spent media (sup) overnight and then subjected to 3% $H_2O_2$ killing for 1h. **(B)** USA300 and *crtM*::Tn were grown overnight in the presence of 50μg/mL flavone (+F) to inhibit STX production, mixed with an equal amount of PAO1, and subjected to 3% $H_2O_2$ killing for 1h. **(C,D)** USA300 and *crtM*::Tn were pre-treated with or without 5% (v/v) filter-sterilized PAO1 or Δ*pqsA* spent media (sup), or 5μM HQNO or PQS overnight, mixed with an equal amount of PAO1, and subjected to 3% $H_2O_2$ killing for 1h in either LB **(C)** or SCFM2 **(D)**. **(E)** USA300 was pre-treated with or without 20% (v/v) filter sterilized PAO1Δ*pqsA*, PDO300 or PDO300Δ*pqsA* spent media (sup), mixed with an equal amount of PDO300, and subjected to 3% $H_2O_2$ killing for 1h. *S. aureus* survival is presented as CFUs normalized to the starting CFUs at 0h. Data presented as mean ± SD from the results of at least 3 biological replicates, each with 3 technical replicates. ns, not significant, compared to USA300 with no treatment (-).

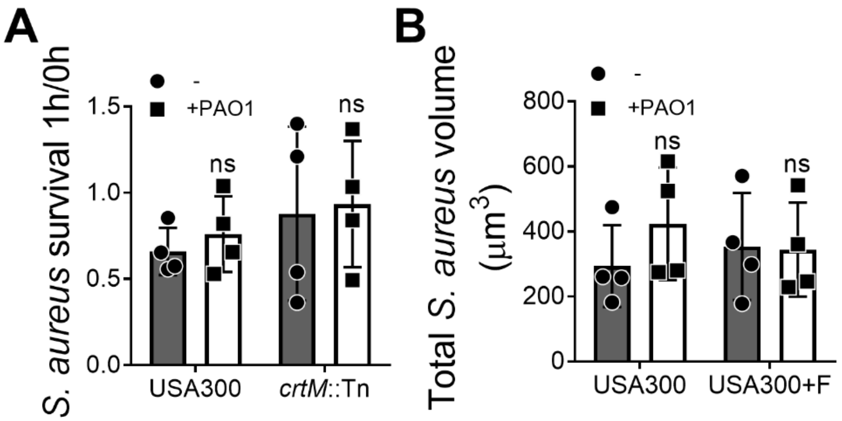


**Figure S5. *S. aureus* survival in the presence of human neutrophils. (A)** USA300 or *crtM*::Tn, either alone or mixed with an equal amount of PAO1, was subjected to human neutrophil killing for 1h (MOI = 10 for each species). *S. aureus* survival is presented as CFUs normalized to the starting CFUs at 0h. Data presented as mean ± SD from the results of at 4 biological replicates, each with 3 technical replicates. **(B)** USA300-GFP was pre-treated with 50μg/mL flavone (+F) to inhibit STX production, then mixed with or without an equal amount of PAO1-TdTomato, was added to adhered human neutrophil (PMN) for 1h. Total *S. aureus* volume was measured by NIS-Element AR software. Data presented as mean ± SD from the results of at least 4 biological replicates, each with 6 technical replicates. ns, not significant, compared to *S. aureus* without the presence of PAO1 (-).

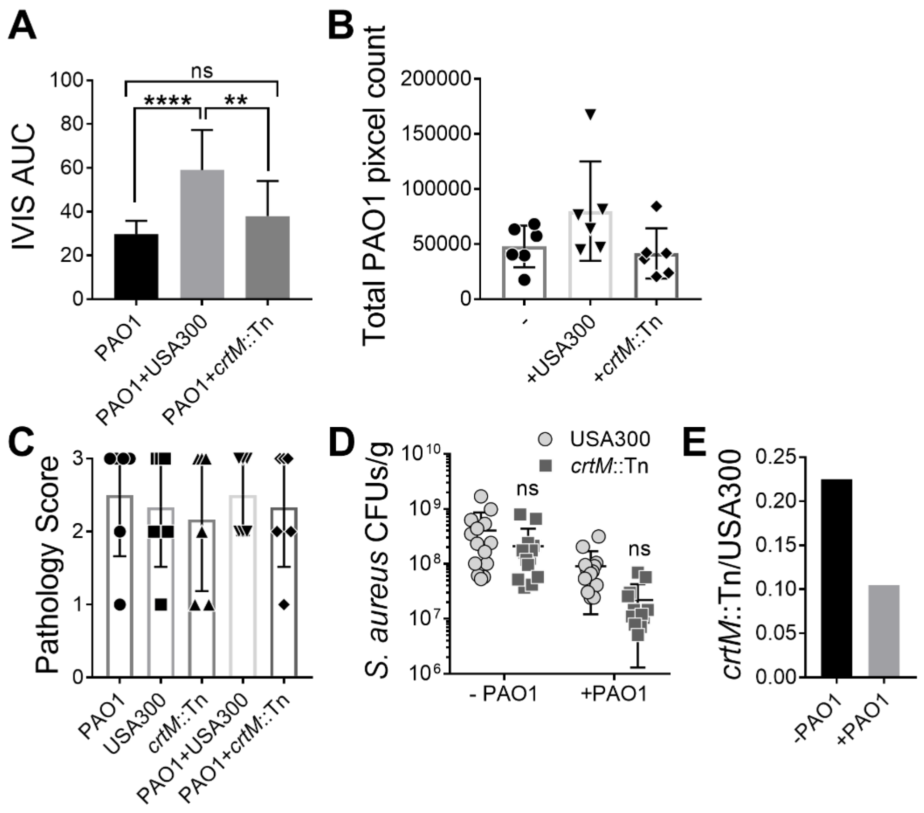


**Figure S6. Analysis of *P. aeruginosa* and *S. aureus* co-infection in the mouse wound model.** **(A)** AUC of Figure 7D comparing PAO1 bioluminescent signal intensity among PAO1 mono-infection and co-infections with USA300 or *crtM*::Tn through the 3-day infection. **, $P < 0.01$; ****, $P < 0.0001$; ns, not significant. **(B)** PAO1 total pixel count from IF-stained wound sections among all groups was quantified. Data presented as mean ± SD from the results of 6 biological replicates. **(C)** Pathology scores of the wound tissues 3 days after infection among all groups. Data presented as mean ± SD from the results of 6 biological replicates. **(D)** USA300 and *crtM*::Tn CFUs/g among all groups were quantified. Data presented as mean ± SD from the results of at least 12 biological replicates, each with 3 technical replicates. ns, not significant, compared to USA300 infection. **(E)** The ratio of *crtM*::Tn survival to that of USA300 was compared between *S. aureus* mono-infection (-PAO1) and co-infection with PAO1 (+PAO1).

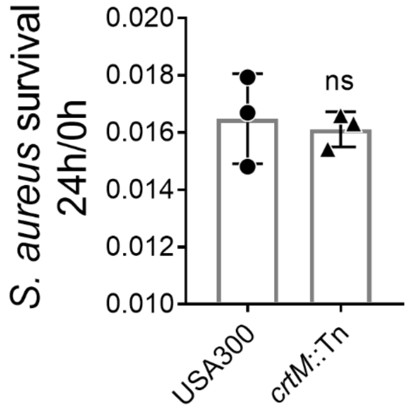


**Figure S7. *S. aureus* survival in co-culture with PAO1.** USA300 or *crtM*::Tn were cultured with PAO1 in LB for 24h. *S. aureus* survival was quantified by comparing CFUs at 24h to that of 0h. Data presented as mean ± SD from the results of 3 biological replicates, each with 2 technical replicates. ns, not significant, compared to USA300.
